# Combining lineage correlations and a small molecule inhibitor to detect circadian control of the cell cycle

**DOI:** 10.1101/2024.03.29.587345

**Authors:** Anjoom Nikhat, Shaon Chakrabarti

## Abstract

Chronotherapy has emerged as an exciting possibility for improving treatment regimens in cancer. A strong influence of the circadian clock on the cell cycle is a crucial requirement for successful chronotherapy. However, though a number of molecular interactions have been discovered between these two oscillators, it remains unclear whether these interactions are sufficient to generate emergent control of cellular proliferation. In this work, we computationally explore a strategy to detect clock control over the cell cycle, by computing lineage correlations in cell cycle times in the presence and absence of the clock inhibitor KL001. Using phenomenological models, we and others have previously suggested that the ‘cousin-mother inequality’ – a phenomenon where cousin cells show stronger cell cycle time correlations than mother-daughter pairs, could be leveraged to probe circadian effects on cellular proliferation. Using stochastic simulations calibrated to match HCT116 colon cancer proliferation datasets, we demonstrate that the established gene-networks giving rise to the cell cycle and circadian oscillations are sufficient to generate the cousin-mother inequality. In the presence of KL001 which stabilizes CRY1, our models predict greater than 50% decrease in the cousin-mother inequality, but counter-intuitively, very little change in population growth rates. Our results predict a range of underlying cell cycle times where the cousin-mother inequality should be observed, and consequently suggest the exciting possibility of combining measurements of lineage correlations with KL001 as a probe of circadian clock – cell cycle interactions.

## Introduction

The idea that cell proliferation exhibits rhythmicity dates back to at least the 1940’s, when Klein and Geisel reported ∼ 24 hour oscillations in the mitotic rate of duodenal epithelium of mice and rats [1]. This remarkable observation suggested that the cell cycle in the alimentary tract of rodents was under the control of the circadian clock, leading to peaks and troughs over time in the fraction of cycling cells. A number of subsequent studies arrived at the same conclusion, though demonstrating that there could be a large variation in the amplitude and mean levels of the mitotic oscillations depending on the particular region of the alimentary canal under study [2, 3, 4, 5]. Further studies extended these results to other tissues such as the skin, bone marrow, tongue and cornea, some of which showed persistent rhythms even under conditions of constant darkness, implying that the rhythmicity was being imposed by the endogenous circadian clock and not the external light-dark cycles [6, 7]. Decades later, these largely physiological studies received strong support when the first molecular links between the circadian clock and cell cycle were discovered. Transcription of two important genes responsible for cell cycle progression – *c-myc* [8] and *Wee1* [9], were found to be directly under the control of circadian clock proteins, via interactions with E-box regions in their promoters. It was also demonstrated, at least in liver hepatocytes, that the control was unidirectional – the clock regulated cell proliferation and not the other way around [9]. Eventually a number of other molecular connections between the two cellular oscillators were also discovered, primarily acting on the G1-S and G2-M transitions of the cell cycle [10, 11, 12]. Taken together these studies provided strong evidence that mitotic oscillations exist, allowing for the exciting prospect of chronotherapy in diseases like cancer, where administering drugs during the peak of fast cycling cancer cells could potentially maximize their eradication [13, 14, 15, 16, 17].

However, the idea of oscillations in cell proliferation and timing therapy to hit the mitotic peaks has consistently been tempered by studies that have failed to find rhythms in cell proliferation. Since the early study on albino rats by Leblond and Stevens that reported a constant renewal of the intestinal epithelium [18], a number of subsequent studies similarly concluded that rhythms in cell proliferation are absent [19, 20]. While this earlier debate was based primarily on measurements of bulk datasets (averages over millions of cells), more recent studies using single cells and time-lapse microscopy have also thrown up contradictory results [12]. Indeed, one such study using a fluorescent reporter of the circadian clock in mouse fibroblasts (NIH3T3 cells) arrived at the conclusion that it is the cell cycle that exerts a strong control over the circadian clock [21], not the other way around as has been conventionally believed [22, 12] and also concluded from recent theoretical studies [23]. This demonstrates the need for developing more accurate approaches to detect the presence of and resolve the directionality of clock-cell cycle coupling, ideally using methods that are robust to the details of the underlying techniques (for example endogenous versus transgenic reporter systems, kinetics of fluorophore maturation [24], and strong assumptions underlying mathematical models).

A potentially exciting method to detect oscillatory control of the cell cycle was recently suggested [25, 26], based upon an observation made in a series of recent studies – many cell types exhibit higher correlations in cell cycle times among cousins as compared to mother-daughter cell pairs (the so called ‘cousin-mother inequality’) [25, 26, 27]. The origin of this puzzling phenomenon was suggested to be an oscillatory control over cell cycle speed [25, 26], and the oscillation time-period that best recapitulated measured correlations and cell cycle time distributions was around 24 hours [27]. The natural candidate for such an oscillatory driver of the cell cycle was the circadian clock, and this expectation was further strengthened when deletion of the clock in bacteria abrogated the lineage correlation structure [26]. Investigating the presence of the cousin-mother inequality was therefore suggested as a potential approach to detecting clock control over the cell cycle [26]. However, the mathematical models originally used to explain the correlations were phenomenological in nature [25, 26, 27], hence it remains unclear whether the established gene-networks underlying the circadian clock and cell cycle oscillators, along with their known molecular couplings, can be expected to generate the experimentally observed correlation structures. Investigating the emergence of lineage correlations from the gene-networks becomes especially relevant since noise in the underlying network dynamics can potentially mask the cousin-mother inequality [26] and also affect the correlation structure in general [28]. Furthermore, recent theoretical studies have shown that oscillatory control over the cell cycle is not the only way to obtain such correlation structures [29, 23]. Therefore simply the presence or absence of the cousin-mother inequality is unlikely to be sufficient to establish the presence of cell cycle control by the circadian clock.

To investigate approaches that can overcome the above-mentioned challenges, here we develop stochastic models to study the emergence of lineage correlations from the known underlying gene-networks of the coupled clock-cell cycle oscillators. Our results demonstrate that in a model calibrated to recapitulate measured noise in HCT116 cells, the ‘forward’ coupling between the clock proteins BMAL1-CLOCK and the cell cycle gene *Wee1* is sufficient to generate the cousin-mother inequality. Additionally, by incorporating a ‘reverse’ coupling (i.e. inhibition of the circadian clock gene *Rev-erb-α* by the CycB/CDK1 cell cycle proteins), we provide the crucial check that clock control by the cell cycle cannot explain the observed lineage correlations. Finally, we model the action of the small molecule inhibitor KL001 which stabilizes the CRY protein, increasing the circadian clock time-period and decreasing the amplitude in a concentration dependent manner [30]. Simulating the effect of KL001 on the clock, we interestingly predict a large percentage decrease in the cousin-mother inequality, as opposed to the proliferation rate which exhibits a much smaller and potentially undetectable decrease. Our results therefore suggest that in addition to the cousin-mother inequality, a large decrease in the inequality upon administration of KL001 is expected to result from the dynamics of the underlying gene-networks. Combining KL001 along with lineage-tracking microscopy could therefore be a simple yet sensitive assay for detecting circadian clock control of the cell cycle.

## Methods

### Models 1 and 2 for the circadian clock and cell cycle gene networks

We model the coupled Circadian clock - cell cycle network schematised in Figure 1A using Ordinary Differential Equations(ODEs), composed of Michaelis-Menten kinetics combined with first and second order reaction kinetics. Owning to the inherent stochasticity of gene expression in biological systems we add Gaussian noise to this model using a Chemical Langevin Equation framework to convert the ODEs into stochastic differential equations (SDEs), and integrate them using the Euler-Maruyama scheme [31]. Details of the models and parameter values used can be found in the SI Sections 1 and 2, and the integration scheme in SI Section 3.

**Figure 1:**
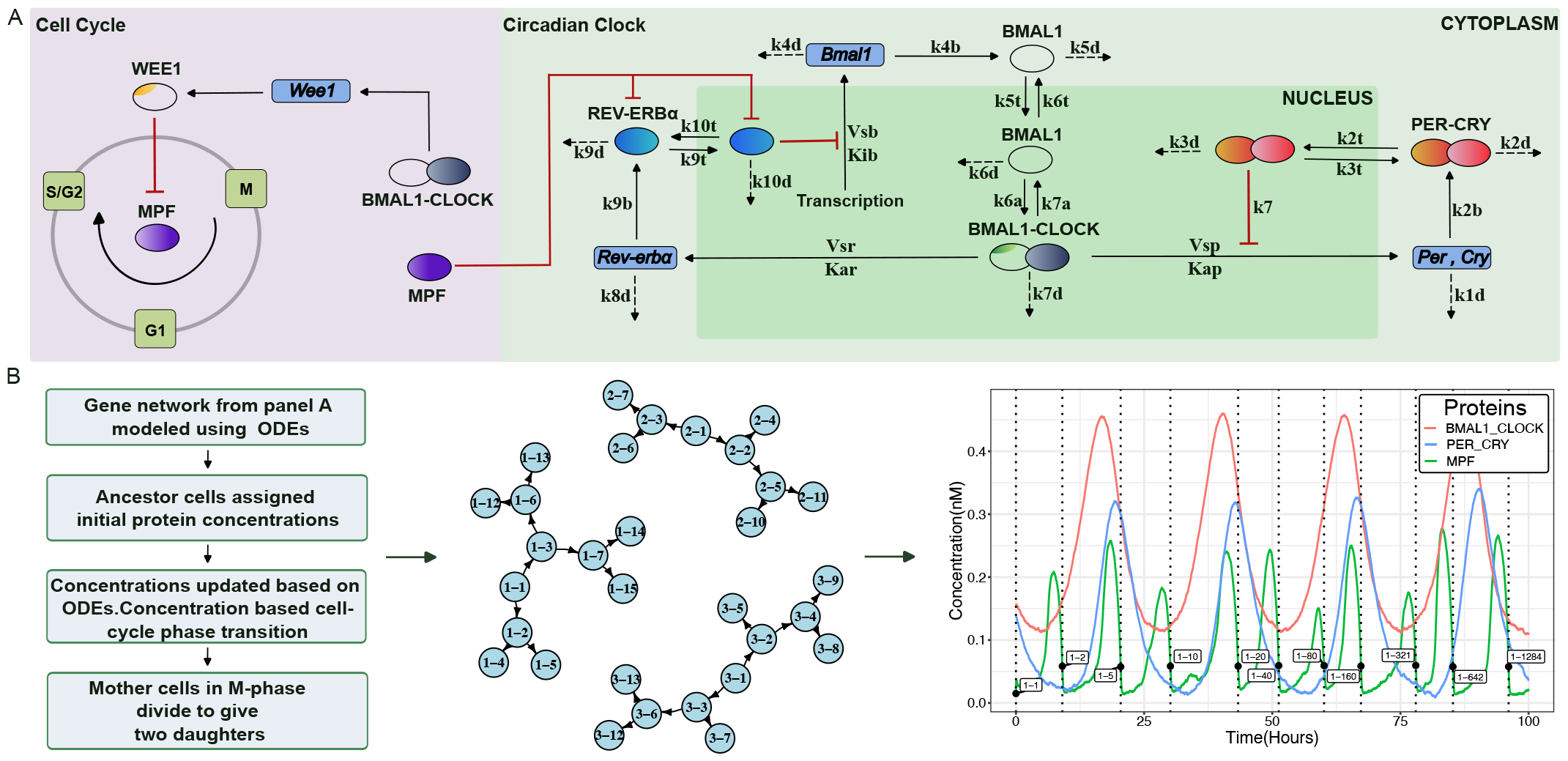
An overview of the coupled circadian clock - cell cycle gene network and the single cell lineage simulation. (A) The coupled circadian clock - cell cycle gene network (Model 2) includes the transcriptional-translational feedback loops that generate the circadian oscillations. The circadian clock gene network shown in the right panel comprises of the auto-regulatory PER-CRY loop along with the REV-ERV*α* mediated oscillatory dynamics of BMAL1. In order to depict the nuclear-cytoplasmic translocation processes we have explicitly separated the nucleus and the cytoplasm in the schematic. The cell cycle as shown in the left panel is divided into three phases-G1, S/G2 and M. Mitosis Promoting Factor (MPF) promotes progression through the phases. Forward and Reverse coupling between the two oscillators are via BMAL1-CLOCK control over *Wee1* and MPF mediated inhibition of REV-ERB*α* respectively. (B) The workflow of our lineage simulation and its output. Left: Algorithm. Middle: A lineage constructed by the simulation where each cell is named as a combination of the lineage it belongs to and the cell number within that lineage at the time of its birth. Right: Time evolution of different proteins for a lineage, recapitulating oscillations with correct phase relationships. The cells belonging to a lineage that were tracked to generate these oscillatory trajectories are labelled at the time of their birth on the graph.

### Lineage simulations to extract correlations in cell cycle times

To study the effect of coupled gene network on the Inter-Mitotic times of related cells, we developed a lineage simulation framework where the clock-cell cycle networks are simulated for individual cells, cell division is driven by the levels of cell cycle proteins, and finally inheritance of all cellular components is symmetric at time of division (details in SI Section 5). The IGRAPH package in R was used to represent the lineage as a directed graph [32, 33]. We initialise a population of ancestral cells and assign the concentration of the different molecular species to each of them. Numerically integrating the system of SDEs drives the cells through cell cycle. Once the mother cells reach the M-phase, we introduce two daughter cells that have the same concentrations as their mother, and remove the mother cell from the population. The simulation runs until a user-defined population size is reached. Once we have the lineage graph we create unique related cell pairs and study the IMT correlation between them.

### Modeling the effect of KL001 on the circadian clock

We incorporate the effect of a small molecule inhibitor of the circadian clock, KL001, which prevents ubiquitin-mediated inhibition of CRY proteins thereby stabilising their concentration. This leads to loss of oscillations in concentration of circadian clock genes [30]. Since we do not explicitly have CRY proteins in our gene network, we simulate KL001’s effect by decreasing the degradation rate of the PER/CRY complex in our model (see SI Section 6 for details). To model increasing concentration of KL001, we divide the PER/CRY degradation rate by certain factors; the factor is set to 1 for the control case where no KL001 is present.

## Results

### Stochastic biochemical models for the coupled circadian clock - cell cycle gene network in single cells

To explore the possibility of lineage correlations arising from the biochemical networks underlying the circadian clock and the cell cycle, we use two models for the coupled oscillators – Model 1 and Model 2, which vary slightly in the number of molecular components involved. The more complex Model 2 is schematically shown in Figure 1A, showing the coupled circadian clock - cell cycle gene regulatory network. We briefly describe the components of the model here; a flowchart of the model simulations is presented in Figure S1 and detailed equations and parameter values are provided in Methods and SI Section 1 and 2. Our circadian clock model was inspired by a more detailed model previously suggested in the literature [34, 35]. We incorporated the negative auto-regulatory loops involving the PER and CRY genes, whose protein products upon translation and translocation back to the nucleus inhibit their own transcription (Model 1). In addition, a secondary loop where the oscillatory dynamics of BMAL1 protein is mediated by REV-ERB*α* was included in the more complex Model 2. BMAL1-CLOCK heterodimer activates the transcription of *Rev-erbα* mRNA, whose protein product is responsible of inhibiting *Bmal1* transcription thereby decreasing active BMAL1-CLOCK concentration as shown in Figure 1B. When the inhibition is lifted due to decreasing concentration of REV-ERB*α*, BMAL1-CLOCK concentration increases thereby restarting the oscillation. Both these models lead to 24-hour oscillations of the mRNA and protein concentrations as shown in Figure 1B and Figures S2.

The cell cycle model used was previously suggested in literature [35]. The mammalian cell cycle is dependent on the activity of multiple Cyclin and Cyclin-Dependent Kinase(CDK) complexes. However in this simplified network, the authors considered the progression through the cell division process to be dependent on the activity of only CyclinB-CDK1 complex or the Mitosis Promoting Factor (MPF). The circadian gating of the cell cycle or forward coupling was incorporated via the control of inhibitory kinase WEE1 by heterodimer BMAL1-CLOCK. As shown in Figure 1A, WEE1 inhibits MPF, keeping its concentration low for the beginning of the cell cycle, till MPF activity rises above WEE1, at which point cell enters the M-phase and ultimately divides. The effect of the cell cycle on the circadian clock or reverse coupling (only present in Model 2) was incorporated via the CDK1 mediated inhibition of REV-ERB*α* [36]. Since we did not explicitly have CDK1 in the cell cycle model, reverse coupling was introduced via REV-ERB*α* inhibition by MPF or CycB/CDK1 complex. The strength of forward and reverse coupling are mediated by changing *C*1 and *C*2 values respectively in the set of ODEs mentioned in SI.

We converted the above reaction schemes into Stochastic Differential Equations and incorporated them into single cell lineage simulations that mimic the cell division process as demonstrated in Figure 1B (see SI Section 1, 2 and 3 for details of the equations and the numerical integration method). In the simulations, the cell cycle was divided into three phases - G1, S/G2 and M. Transition through the various phases was based on the concentration of MPF, and the ones that ultimately reached the M-phase divided to give two daughter cells. Cellular lineages were represented using directed graphs, where the relationship between the cells was preserved, to enable inference of lineage-relationships between cell pairs for downstream analysis (details in SI Section 5). As shown in Figure 1B, the concentrations over time of different proteins for an extracted lineage recapitulated 24 hour oscillations, where the anti-phase expression of BMAL1/CLOCK complex and the PER/CRY complex could be observed as expected from previous experiments [37].

### Circadian clock mediated forward coupling recapitulates experimentally observed correlation structure in cellular lineages

With the models for the two oscillators and correct phase relationships between the trajectories of MPF, BMAL-CLOCK and Wee1 established, we next asked whether our models exhibit entrainment, as expected from the theory of coupled oscillators. Keeping the circadian clock period (∼ 24 hours) and the autonomous cell cycle period (∼ 20 hours) fixed, we either increased the forward or the reverse coupling strength. We simulated the time evolution of the gene network and took Fourier Transforms to identify the frequency of oscillations of both oscillators (see SI section 4 for details). As shown in Figure 2A, the frequency of oscillation of the cell cycle protein MPF changes from 1/20 hour^*−*1^ when coupling is 0 to 1/24 hour^*−*1^ in presence of high forward coupling, suggesting entrainment of the cell cycle by the circadian clock. Similarly, in Figure 2B, the frequency of BMAL1/CLOCK oscillation shifts to 1/20 from 1/24 hour^*−*1^ with the increase in reverse coupling strength, reflecting entrainment of the circadian clock by the cell cycle.

**Figure 2:**
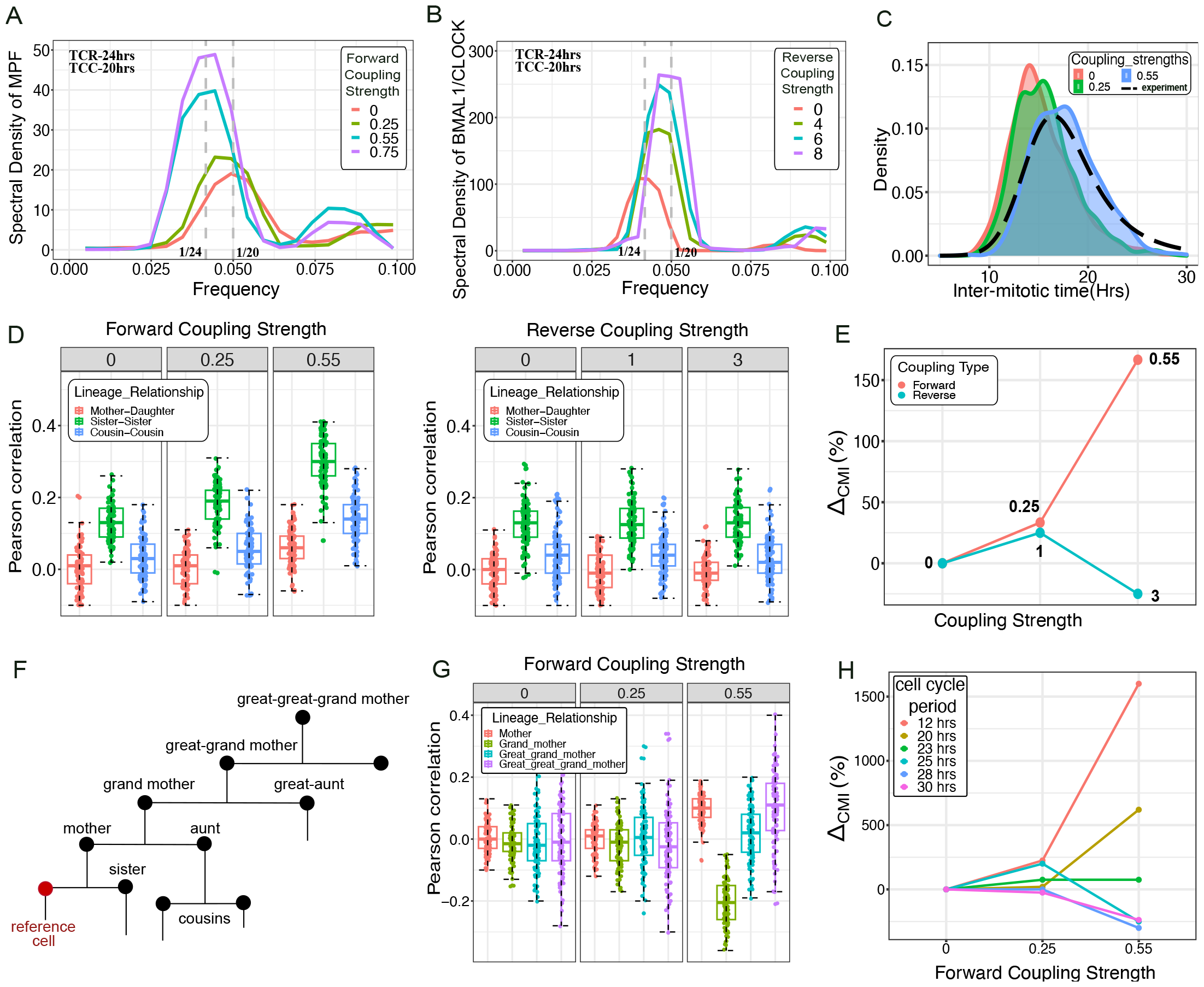
Circadian clock control over the cell cycle via BMAL1-CLOCK interaction with *Wee1* is sufficient to generate the cousin-mother inequality. (A) Entrainment of the cell cycle by the circadian clock. With increasing forward coupling strength the frequency of oscillation of the cell cycle gene MPF changes from 1/20 to 1/24 hour^*−*1^. (B) Entrainment of circadian clock by the cell cycle. With increasing reverse coupling strength the circadian clock heterodimer BMAL1/CLOCK oscillation frequency changes from 1/24 to 1/20 hour^*−*1^. Circadian clock period (TCR)=24 hrs, Autonomous cell cycle period (TCC)=20 hrs. (C) Comparison of IMT distribution from simulation and experiment. The simulated IMT distribution (with coupling strength 0.55) approximately matches the experimentally observed IMT distribution (black dashed line). (D) Increasing forward coupling strength generates high sister correlations and also gives rise to the cousin-mother inequality. In contrast, reverse coupled system fails to recapitulate the cousin-mother inequality. Boxplots were generated from 100 runs of the simulations. (E) Percentage change in the difference between median values of cousins and mother-daughter correlation in comparison to the uncoupled system, shown for the different coupling strengths. Numbers in the plot refer to the coupling strength either for forward (red) or reverse (blue) coupling. (F) Schematic to demonstrate the lineage relationship considered with respect to the reference cell marked in red. (G) Comparison of the IMT correlation down a vertical generation for the different forward coupling strengths. We observe the negative grandmother correlation (green) as seen in the experiments on HCT116 cells. (H) Effect of different autonomous cell cycle periods on the cousin-mother inequality. Our model predicts *Wee1* mediated forward coupling gives rise to cousin mother inequality when the average cell cycle period is ∼ 12 to 24 hrs.

Having demonstrated that our coupled oscillator model generates the expected entrainment behaviour, we next asked if the forward and reverse coupling can generate lineage related intermitotic time (IMT) correlation structures observed in previous experiments [27]. In particular, we and others have shown that cousin cells often exhibit stronger IMT correlations than mother-daughter pairs (the cousin-mother inequality), and that this surprising phenomenon may arise from circadian gating of the cell cycle [25, 26, 27]. To study the origin of the cousin-mother inequality, we utilised our previously discussed lineage simulation that creates ancestor cells’ lineages based on the stochastic gene network provided as input.

Since noise in the cell cycle times can affect the correlations [27], we determined the optimum noise level and coupling strength for our model by comparing the IMT distribution generated by our simulation to that observed experimentally in the HCT116 cell line [27]. As shown in the blue distribution in Figure 2C and in Figure S3, we were able to approximately reproduce the experimentally observed IMT distribution both for a forward and reverse coupled gene network respectively. We then extracted related cell pairs from our simulations, and compared their cell cycle time correlations for both forward and reverse coupled cases separately. We observed the cousin-mother inequality emerge upon increasing the forward coupling strength, showing a significant increase in the cousin correlations at the coupling strength of 0.55 where the IMT distribution matched experimental results (Figures 2D, E). In contrast, in the reverse coupled gene network where the cell cycle influences the circadian clock, the cousin-mother inequality was not observed even for a high reverse coupling strength (Figures 2D, E). Additionally, to check whether having different levels of noise for the clock and cell cycle networks affects the observation of the cousinmother inequality, we ran a set of simulations with varying degrees of noise strength for the circadian clock components. We found that the inequality was always observed (Figure S3).

We also extracted the vertical generation (grandmother, great-grandmother, etc) correlations in the cell cycle times (Figure 2F, G). As was observed in the experiment [27, 23], the grandmother correlation was negative, thereby suggesting that our model results are consistent with the overall lineage correlation structure measured using time lapse imaging of HCT116 cells.

Finally, we also explored the effect of changing the autonomous cell cycle time on the lineage correlations. Since mammalian cells can have both shorter (∼ 12 hours for stem cells) as well as larger (∼ 24 hours for fibroblasts) average cell cycle times, we varied the mean of the IMT distribution within 12 − 30 hours. The results of these simulations (Figure 2H) predict that the cousin-mother inequality will be stronger at lower IMT values, and is lost closer to average cell cycle times of ∼ 24 − 30 hours. Interestingly, this is seemingly consistent with measurements on mouse fibroblasts (average division time ∼ 24 hours) where the inequality was not observed [38], and HCT116, L1210 cells (average division time of both cell types ∼ 16 hours) where a strong cousin-mother inequality was observed [27, 25]. Note however that whether a forward coupling exists in NIH3T3 cells, is currently debated [21].

Therefore in summary, our results in this section suggest that circadian clock control over the cell cycle (via BMAL1-CLOCK interactions with *Wee1*) is expected to give rise to the cousin-mother inequality, but only when the average autonomous cell cycle time is approximately within 12 − 24 hours.

### KL001 mediated inhibition of the circadian clock leads to a large decrease in the cousin-mother inequality

Though our results from the last section demonstrated that the molecular coupling between BMAL1-CLOCK and *Wee1* can generate the cousin-mother inequality, observation of this inequality in a dataset does not necessarily prove the existence of circadian clock – cell cycle coupling. Recent studies suggested that imposing a cell size dependent regulation of cell cycle speed, can also give rise to similar IMT correlations [29, 23]. Therefore we next asked if perturbations to the circadian clock, and hence the lineage correlations, could potentially be a way to probe the existence of forward coupling between the clock and the cell cycle. Selectively modulating the circadian clock can be achieved experimentally using a multitude of small molecules, such as KL001 and SR9009. KL001 for example stabilizes the CRY protein by preventing its ubiquitin mediated degradation leading to damped oscillations with decreased amplitude and increased time period [30]. A particularly attractive feature of KL001 is that its effects are concentration dependent [30]. We therefore modeled KL001 in our lineage simulations, to study its effects on the correlation structure in cell cycle times.

Since we did not explicitly have CRY proteins in our model, we mimicked the effect of KL001 by decreasing the degradation rates of the PER-CRY complex, which leads to damped circadian oscillations as shown in Figure 3A, similar to ones experimentally observed in [30]. We divided the degradation rates of the PER-CRY complex by the indicated number to simulate addition of increasing concentration of KL001. Thus, the un-inhibited or control system is depicted when KL001 value is equal to 1 (see details in SI section 6). To carefully compare and contrast the experimental results with our simulations, we digitally extracted KL001 time-series datasets from [30] (Figure 3B, C), and computed the time periods and amplitudes of the oscillations as a function of KL001 concentration (Figure 3B, C; see details in SI Section 7). The experimental data showed an abrupt jump in the time period on increasing KL001 concentration, which was well captured by our model (Figure 3B, D). The amplitude on the other hand showed a more continuous decrease, which was also recapitulated in our simulations (Figure 3C, E). The only aspect of the experimental data that did not precisely match our simulations, was a remnant oscillation even at very high KL001 concentrations, resulting in a non-monotonic behavior of the measured time period (Figure 3B). However, this difference is unlikely to affect our results, since at these high concentrations of KL001 the amplitude of oscillations was essentially zero (Figure 3C, E). Overall, our KL001 model therefore seems to largely capture the observed effects of the inhibitor on clock oscillations.

**Figure 3:**
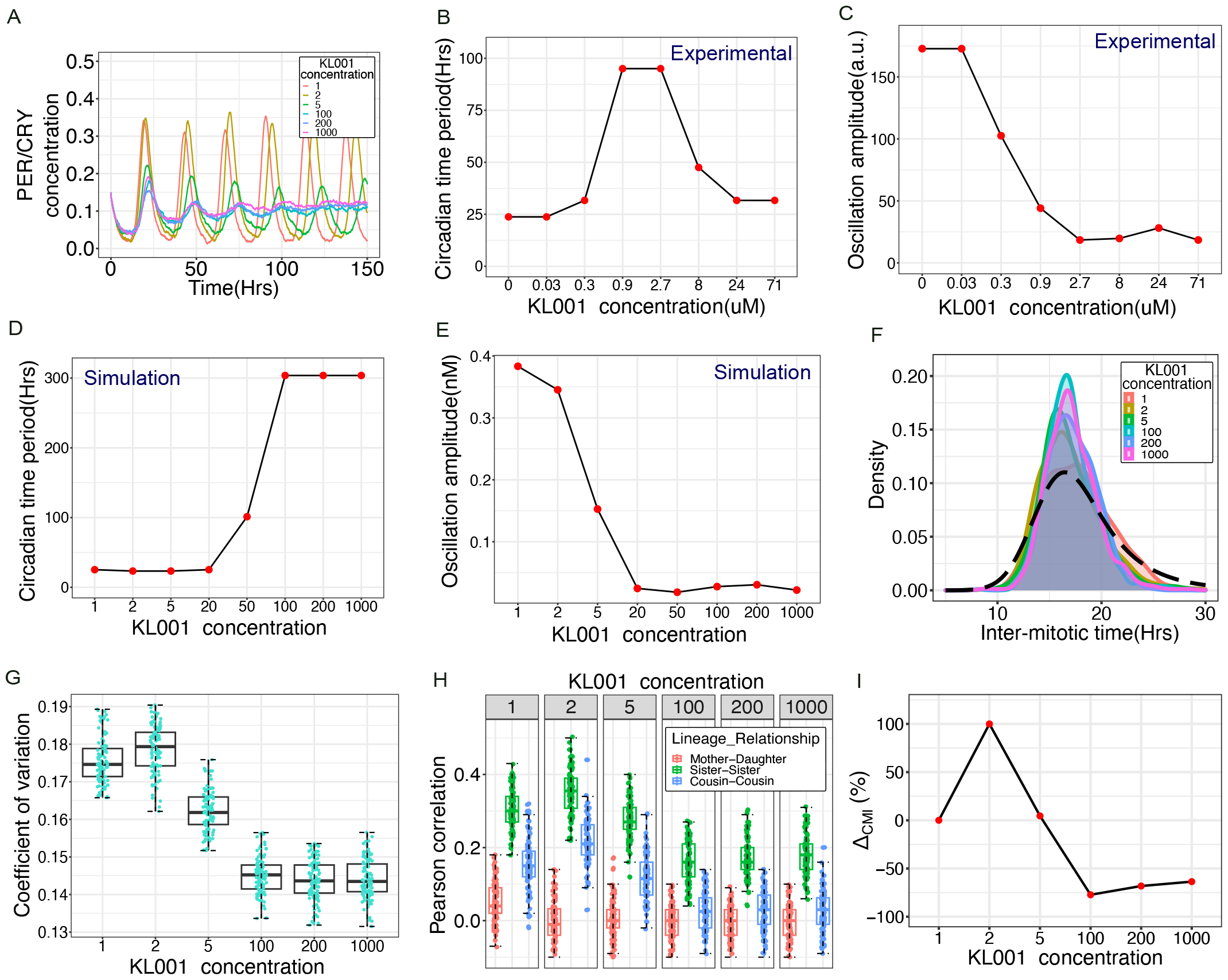
Circadian clock perturbation with KL001 decreases IMT correlations in related cell pairs. (A) Damped circadian clock oscillations as observed with increasing KL001 inhibitor concentration in our Model 2 simulations. (B)&(C) Oscillation time period and amplitude respectively as functions of increasing KL001 concentration. Time series datasets were digitally extracted from experimental results in [30] and analyzed as described in Methods and SI Section 5. (D)&(E) Oscillation time period and amplitude respectively as functions of increasing KL001 concentration from our Model 2 simulations. (F) Change in Inter-mitotic time distribution as observed with increasing KL001 concentration. The model predicts a decrease in variance of the distribution in comparison to the experimental observation (black dashed line). (G) Decrease in the coefficient of variation of Intermitotic time distribution. (H) Increasing concentration of circadian clock inhibitor KL001 diminishes the cousin-mother inequality. (I) Percentage change in median value of cousin-mother inequality in comparison to the control case, shown for the different concentrations of KL001.(Boxplot and median value calculated for 100 runs of simulation. Here C1 = 0.55, C2=0, TCC=15hrs)

With the model for KL001 established, we next asked how increasing KL001 concentration (by reducing the degradation rate of PER-CRY complex) would affect the lineage correlations. Simulating lineages with circadian clock inhibition by KL001 showed a decrease in variability of distribution of cell-cycle times (Figure 3F), which was further quantified by comparing the coefficient of variation (COV) of the IMT distribution for 100 runs of the simulation as shown in Figure 3G. Similar observations were previously reported in *Cyanobacteria*, where time lapse experiments performed on wild-type and clock deleted mutants revealed decreased cell cycle duration variability in case of the mutant [26]. This reduction in variance was attributed to the loss of clock oscillations in the mutant, which otherwise generates larger variability in cell cycle times. Further, we observe that simulating lineages keeping the forward coupling strength high, with increasing circadian inhibition causes the related cell correlation to decrease at highest inhibitor concentration. When comparing the percentage difference in the median value of cousins and the mother-daughter correlations for the inhibited systems (KL001 = 2 to 1000 represents increasing concentration) in comparison to the control system (KL001 = 1), the cousin-mother inequality diminishes by ∼ 50% (Figure 3H, I).

In summary, our results suggest that the effect of the small molecule clock inhibitor KL001 on clock oscillations can be recapitulated using a relatively simple network model of the circadian clock. Furthermore, addition of KL001 at high concentrations is expected to significantly reduce the cousin-mother inequality in cell cycle time correlations, that should be detectable using live cell microscopy techniques.

### Population growth rates are relatively unaffected by KL001-induced clock inhibition

A natural, and in principle simpler alternative to measuring KL001-induced changes in the lineage correlations would be to measure cell proliferation rate changes. Inhibiting the clock oscillations, and thereby the coupling to the cell cycle, would be expected to also change the proliferation rate [35]. As we demonstrated earlier in Figures 2A-B, entrainment causes the average cell cycle time to shift closer to the clock period, hence one might expect KL001 treatment to revert the clock-driven proliferation rate back to the autonomous (no coupling) scenario. If this were to be the case, singlecell lineage tracking experiments would not be required, and simply measuring population dynamics in bulk assays would provide an easier approach to detecting clock control of the cell cycle.

Using our lineage simulation, we generated trajectories of cell population as a function of time as shown in Figure 4A. We fit a linear model to the *log* (cell number) *versus* time data, to estimate the proliferation rate of the population. First we checked our model results for increasing forward coupling strength, and found that the proliferation rate decreased as expected due to entrainment (Figure 4B). However, under circadian clock inhibition with KL001, the proliferation rate interestingly showed minimal increase of ∼2% (Figure 4C). This result can be explained by observing that the mean of the cell cycle times distribution does not change with increasing KL001 concentration (Figure 3F).

**Figure 4:**
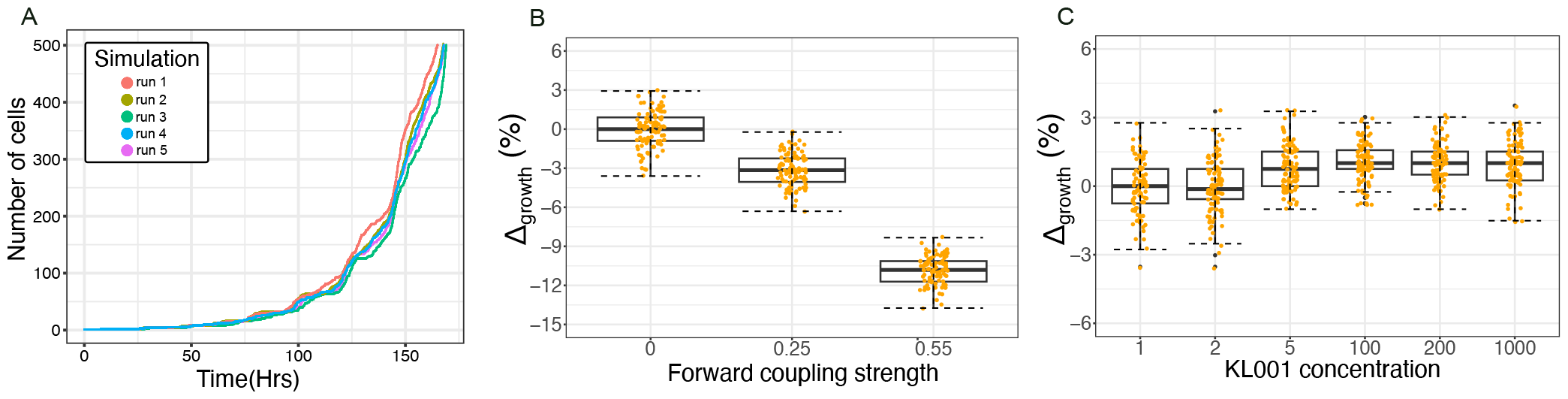
Cell proliferation rates are predicted to remain relatively unaffected by circadian clock inhibition using KL001. (A) Representative exponential trajectories of number of cells in a population as a function of time for different runs of the simulation. (B) Decreasing proliferation rate observed for increasing forward coupling strength owing to entrainment of the cell cycle by the circadian clock. We plot the percentage change in the median proliferation rate relative to the uncoupled system i.e. *C*1 = 0. (C) The cell proliferation rate remain unaffected with increasing KL001 mediated circadian clock inhibition. Here the proliferation rate is plotted as percentage change relative to the uninhibited system depicted by KL001 = 1. (Boxplot and median value calculated for 100 runs of simulation. Here C1 = 0.55, C2=0, TCC=15hrs)

Our results in this section predict that the proliferation rate increase upon adding KL001 to proliferating cells will be minimal, and likely undetectable within experimental resolution. Changes in the lineage correlations are much larger as we found in the previous section, and hence will likely be a more powerful approach to detecting circadian control over the cell cycle.

## Discussion

One of the key ideas behind cancer chronotherapy is administration of treatments at a time of day when cancer cell proliferation is at its peak [39]. The important assumption underlying this idea is that circadian clock control over the cell cycle will lead to oscillations over time in the fraction of cells in any cell cycle phase [40, 17]. However, even though substantial evidence exists for a molecular level coupling of the clock and the cell cycle, whether there is any emergent control of cell proliferation by the clock remains a heavily debated subject. The question of emergent control is particularly relevant since the circadian clock oscillations are known to be low-amplitude and noisy, and therefore molecular interactions with the cell cycle are not guaranteed to result in strong control of proliferation. The key unanswered questions therefore are whether the underlying microscopic dynamics at the level of the proteins and mRNA in single cells is sufficient to generate cell cycle control by the oscillating circadian clock, and how to establish sensitive approaches to demonstrate evidence for such control, if it exists.

Here we use stochastic, mechanistic models to predict that one of the best studied molecular connections – BMAL1-CLOCK driving of the cell cycle via the G2-M inhibitor *Wee1*, is likely to be sufficient to exert detectable control over cellular proliferation as measured by lineage correlations in cell cycle times. By carefully calibrating the noise and coupling coefficients, we ensure that our models generate distributions of cell cycle times that are similar to those measured experimentally in HCT116 cells. Given that these distributions can strongly affect the lineage correlations [26, 27, 28, 38], our results demonstrate that realistic noisy dynamics of mRNA and proteins can still generate the cousin-mother inequality in cell cycle time correlations. Perhaps more importantly, we show that our simulated KL001 treatment, which recapitulates the experimentally observed abrupt increase in clock time period and more continuous reduction in amplitude, gives rise to a large reduction (*>* 50%) in the cousin-mother inequality. Interestingly, such a large change is not observed in the population growth rate of the cells, since even very high KL001 concentrations only flatten out the PER-CRY/BMAL1-CLOCK oscillations without reducing their absolute levels to zero (thereby maintaining a steady coupling to the cell cycle). Our results therefore suggest that measuring KL001 mediated reduction in the cousin-mother inequality, but not the population growth rates, could be a sensitive readout for clock control over the cell cycle.

The advantage of our proposed method is that it relies on two relatively simple experiments requiring just the ability to track cell divisions – (i) measuring lineage correlations of cell cycle times *in vitro*, and (ii) repeating the same measurements in the presence of the small molecule inhibitor of the clock, KL001. The method also requires no addition of synchronizing agents such as dexamethasone, but rests on the assumption that the clock phase gets inherited by daughter cells at time of division, thereby maintaining some degree of phase synchrony within lineages [27]. However, we anticipate potential limitations which might render this approach applicable only in limited cell types. For conclusively determining clock control of the cell cycle, our approach requires positive results from both the above experiments – a strong cousin-mother inequality to begin with, and subsequently a large decrease in the inequality upon KL001 addition. A negative result in either experiment is likely to be hard to interpret: the absence of the inequality may arise from particular combinations of cell cycle and clock periods (as we show in Figure 2H) or even simply due to masking by noise [26]. Maintenance of the inequality even in the presence of KL001 might indicate the presence of other stronger hidden variables (such as cell size [29, 23]) that couple to the cell cycle and mask the effect of clock-generated correlations. Finally, there are a few limitations with our modeling approach as well – as with any mechanistic model based on biomolecular reactions, the results may be somewhat dependent on the precise network connections chosen. For example, while the BMAL1-CLOCK connection to *Wee1* is best characterized, other connections between the clock and cell cycle also exist and have been computationally modeled, which we have ignored [41]. Additionally, our models cannot recapitulate the absolute values of the lineage correlations as seen in experiments, but rather explain only the qualitative structure of the cousin-mother inequality. This is expected, since it is well understood that a wide variety of inherited components from mother to daughter cells such as mRNA/protein levels, transcription rate, and epigenetic states could all contribute to setting the absolute values of the correlations [27, 28, 42, 43]. However, since our method conceptually rests upon identifying the inequality in correlation patterns, our inability to reproduce the absolute correlations is unlikely to be a major limitation.

## Conclusions

Recent discoveries of small molecule inhibitors and activators of the circadian clock provide the exciting possibility of manipulating cellular processes strongly coupled to the clock. The cell cycle is one such important potential target, that might be indirectly controlled via perturbations of the clock for chronotherapeutic purposes. Using stochastic computational models, here we provide a novel suggestion of combining measurements of cell cycle time correlations on lineages with the clock inhibitor KL001, to detect whether the clock controls cell proliferation – a question that remains heavily debated in the field. Whether our predictions hold and the conclusions generalizable to a variety of cell types remain to be demonstrated with carefully executed future experiments.

## Acknowledgements

S.C. acknowledges funding from SERB (Government of India) under project number SPR/2021/000486 as well as intramural funds from National Center for Biological Sciences–Tata Institute of Fundamental Research (NCBS-TIFR).

## Author contributions

S.C. conceptualized and designed the study, A.N. developed the models and performed all calculations and analyses, A.N. and S.C. wrote the paper, S.C. supervised the work.

