## Supplementary Information for "Combining lineage correlations and a small molecule inhibitor to detect circadian control of the cell cycle"

March 29, 2024

### Contents

**1 Model 1: A simplified gene network model of circadian clock - cell cycle coupling 2**

**2 Model 2: Description of the coupled circadian clock - cell cycle model 4**

2.1 Simplified Circadian clock model . . . . . . . . . . . . . . . . . . . . . . . . . . . . . . 4

2.2 Coupling the circadian clock - cell cycle system . . . . . . . . . . . . . . . . . . . . . . . 5

**3 Numerical Integration of the system of ODEs to generate time series 7**

**4 Fourier Analysis of time series data 7**

**5 Simulation to generate cellular lineages 7**

**6 Simulating KL001 mediated circadian clock inhibition 11**

**7 Digital extraction of KL001 data and downstream processing 13**

### 1 Model 1: A simplified gene network model of circadian clock - cell cycle coupling

Since only phenomenological models have so far been used to explain the emergence of the cousin-mother inequality in cell cycle time correlations, here we explored whether a very simple network model of the circadian clock coupled to the cell cycle [1] is sufficient to generate lineage correlations. The circadian clock network used in [1], which we describe as Model 1, consists only of the PER-CRY regulatory loop. Model 1 includes the BMAL1-CLOCK mediated activation of transcription of the *Per* and *Cry* mRNAs. *Per* or *Cry* isoforms are not explicitly considered, but instead clubbed together as a single species. The mRNAs are translated in the cytoplasm and upon translocation back to the nucleus they inhibit their own production. This system is represented by a set of 7 coupled ODEs (Equations 1-7).

The mammalian cell cycle is driven by multiple Cyclin and Cyclin-dependent Kinases. In this simplified model the authors however consider the entire cell cycle to be dependent on the concentration of CyclinB-Cdk1 complex or the Mitosis Promoting Factor(MPF) [1]. Progression through each of these phases is dependent on certain concentration thresholds of MPF. The cells in G1 phase enter the S-phase when [MPF] $>$ 0.09 nM. The model also includes a tyrosine kinase WEE1 which inhibits Cdk1 and prevents the G2/M transition. Hence for cells to enter the M-phase the [MPF] $>$ [WEE1]. Once [MPF] surpasses [WEE1] another inhibitor of MPF is activated which is analogous to the Anaphase Promoting Complex (APC). This inhibitor brings the [MPF] down to a lower threshold ($\leq$ 0.06 nM) for the cell to exit mitosis and enter G1 phase. This system is represented by a set of 3 coupled ODEs (Equations 8-10).

The circadian clock mediated forward coupling was included via BMAL1-CLOCk mediated control of kinase protein WEE1 of the cell cycle, thus resulting in control of G2/M transition by the clock. The strength of this coupling is controlled by the value of the coupling constant C1. The variable names corresponding to each species in Model 1 are mentioned in Table S1. The variables associated with circadian clock are described by $y_i$, $i = 1, 2, ..., 7$ and those associated with the cell cycle are indicated by $z_j$, $j = 8, 9, 10$. The parameter values for Model 1 are mentioned in Tables S2 and S3. We numerically integrated the Chemical Langevin form [2] of the set of 10 ordinary differential equations that constituted the model as described later in Section 3.

The control of the autonomous cell cycle period (TCC) in this model was imposed by multiplying the RHS of the 3 cell cycle ODEs by a scaling factor of (97.4/TCC). For Model 1, TCC = 16 hrs.

| Variable | Molecular species |
|---|---|
| $y_1$ | PER-CRY mRNA |
| $y_2$ | PER-CRY cytoplasmic protein |
| $y_3$ | PER-CRY nuclear protein |
| $y_4$ | BMAL1 mRNA |
| $y_5$ | BMAL1 cytoplasmic protein |
| $y_6$ | BMAL1 nuclear protein |
| $y_7$ | BMAL1-CLOCK complex |
| $z_8$ | Mitosis Promoting Factor (MPF) |
| $z_9$ | WEE1 |
| $z_{10}$ | MPF-inhibitor (APC) |

Table S1: Molecular species and corresponding variable names for Model 1.

$$\frac{dy_1}{dt} = \frac{v_{1b}.(y_7 + c)}{k_{1b}.(1 + (\frac{y_3}{k_{1i}})^p) + (y_7 + c)} - k_{1d}.y_1 \tag{1}$$

$$\frac{dy_2}{dt} = k_{2b}.{y_1}^q - k_{2t}.y_2 + k_{3t}.y_3 - k_{2d}.y_2 \tag{2}$$

$$\frac{dy_3}{dt} = k_{2t}.y_2 - k_{3t}.y_3 - k_{3d}.y_3 \tag{3}$$

$$\frac{dy_4}{dt} = \frac{v_{4b}.y_3^r}{k_{4b}^r + y_3^r} - k_{4d}.y_4 \tag{4}$$

$$\frac{dy_5}{dt} = k_{5b}.y_4 - k_{5t}.y_5 + k_{6t}.y_6 - k_{5d}.y_5 \tag{5}$$

$$\frac{dy_6}{dt} = k_{5t}.y_5 - k_{6t}.y_6 + k_{7a}.y_7 - k_{6a}.y_6 - k_{6d}.y_6 \tag{6}$$

$$\frac{dy_7}{dt} = k_{6a}.y_6 - k_{7a}.y_7 - k_{7d}.y_7 \tag{7}$$

$$\frac{dz_8}{dt} = (\frac{k_{0mpf}.k_{1mpf}^{n1}}{k_{1mpf}^{n1} + z_8^{n1} + s.z_{10}^{n1}}).(1 - z_8) - d_{wee1}.z_9.z_8 \tag{8}$$

$$\frac{dz_9}{dt} = (\frac{k_{actw}}{k_{actw} + d_{w1}}).(c_w + C_1.(y_7 - 0.9629)) + (\frac{k_{actw}}{k_{actw} + d_{w1}} - 1).(\frac{k_{inactw}.z_8^{n1}.z_9}{k_{1wee1}^{n1} + z_8^{n1}}) - d_{w2}.z_9 \tag{9}$$

$$\frac{dz_{10}}{dt} = k_{act}.(z_8 - z_{10}) \tag{10}$$

| Parameter Symbol | Numerical Value |
|---|---|
| $c$ | $0.01nM$ |
| $p$ | 8 |
| $v_{1b}$ | $9nMh^{-1}$ |
| $k_{1b}$ | $1nM$ |
| $k_{1d}$ | $0.12h^{-1}$ |
| $k_{1i}$ | $0.56nM$ |
| $k_{2b}$ | $0.3nM^{-1}h^{-1}$ |
| $k_{2d}$ | $0.05h^{-1}$ |
| $k_{2t}$ | $0.24h^{-1}$ |
| $k_{3t}$ | $0.02h^{-1}$ |
| $q$ | 2 |
| $k_{3d}$ | $0.12h^{-1}$ |
| $v_{4b}$ | $3.6nM^{-1}h^{-1}$ |
| $r$ | 3 |
| $k_{4b}$ | $2.16nM$ |

| Parameter Symbol | Numerical Value |
|---|---|
| $k_{4d}$ | $0.75h^{-1}$ |
| $k_{5b}$ | $0.24h^{-1}$ |
| $k_{5d}$ | $0.06h^{-1}$ |
| $k_{5t}$ | $0.45h^{-1}$ |
| $k_{6t}$ | $0.06h^{-1}$ |
| $k_{6d}$ | $0.12h^{-1}$ |
| $k_{6a}$ | $0.09h^{-1}$ |
| $k_{7a}$ | $0.003h^{-1}$ |
| $k_{7d}$ | $0.09h^{-1}$ |

Table S2: Circadian clock ODE parameters for Model 1.

| Parameter Symbol | Numerical Value |
|---|---|
| $k_{0mpf}$ | $10h^{-1}$ |
| $k_{1mpf}$ | $0.05nM$ |
| $n1$ | 2 |
| $s$ | $50nM$ |
| $d_{wee1}$ | $5h^{-1}$ |
| $k_{actw}$ | $1h^{-1}$ |
| $d_{w1}$ | $1nM$ |
| $k_{inactw}$ | $200h^{-1}$ |
| $k_{1wee1}$ | $0.5nM$ |
| $d_{w2}$ | $1h^{-1}$ |
| $k_{act}$ | $0.01h^{-1}$ |
| $c_w$ | 1.46 |

Table S3: Cell Cycle ODE parameter values for both Model 1 and Model 2.

### 2 Model 2: Description of the coupled circadian clock - cell cycle model

#### 2.1 Simplified Circadian clock model

Our circadian clock network in Model 1 only included the PER-CRY mediated negative regulatory loop as mentioned in Section 1, allowing introduction of a forward coupling from the circadian clock to the cell cycle. To allow the reverse interaction, i.e. from the cell cycle to the circadian clock, we introduced a second loop in the circadian clock network where the BMAL1-CLOCK complex also activates the production of *Rev-erb*$\alpha$ mRNA (Model 2). The REV-ERB$\alpha$ protein on translocation back to the nucleus gives rise to oscillations in the concentration of the BMAL1 protein. Decreasing concentration of both PER-CRY complex and the REV-ERB$\alpha$ protein lifts the inhibition causing active BMAL1-CLOCK complex to accumulate, thus resetting the cycle. The modified circadian clock in Model 2 is described by 10 ODEs as shown below (Equations 11 to 20).

The cell cycle network was the same for Model 1 and Model 2 (Equations 8-10), hence the thresholds used to transition cells through the various cell cycle stages remain the same.

#### 2.2 Coupling the circadian clock - cell cycle system

The circadian clock mediated forward coupling is as described before, via the control of cell cycle protein WEE1 by which the circadian clock prevents the G2/M transition [3]. To incorporate the reverse control of the cell cycle on the circadian clock, we incorporated Cdk1 mediated inhibition of REV-ERB$\alpha$ [4]. Since we do not explicitly have Cdk1 in our cell cycle system, MPF was used as the inhibitory link. The strengths of the forward and the reverse couplings are controlled by the coupling constants $C_1$ and $C_2$ respectively in the equations for WEE1 and REV-ERB$\alpha$ proteins.

The reactions were modelled using a combination of Michaelis-Menten kinetics, along with first and second order reaction kinetics. The 10 ODEs for the circadian clock model along with the 3 ODEs describing the cell cycle as in Section 1, together depict the time evolution of the concentrations of different molecular species included as shown in Figure 1A in the main text. The variable names of the species in Model 2 are mentioned in Table S4 and the parameter values for the system of 10 ODEs is provided in Table S5.

| Variable | Molecular species |
|---|---|
| $y_1$ | PER-CRY mRNA |
| $y_2$ | PER-CRY cytoplasmic protein |
| $y_3$ | PER-CRY nuclear protein |
| $y_4$ | BMAL1 mRNA |
| $y_5$ | BMAL1 cytoplasmic protein |
| $y_6$ | BMAL1 nuclear protein |
| $y_7$ | BMAL1-CLOCK complex |
| $y_8$ | REV-ERB$\alpha$ mRNA |
| $y_9$ | REV-ERB$\alpha$ cytoplasmic protein |
| $y_{10}$ | REV-ERB$\alpha$ nuclear protein |

Table S4: Molecular species and corresponding variable names for the Circadian clock network in Model 2.

$$\frac{dy_1}{dt} = v_{sp}.(\frac{y_7^n}{k_{ap}^n + y_7^n}) - k_{1d}.y_1 \tag{11}$$

$$\frac{dy_2}{dt} = k_{2b}.{y_1}^q - k_{2t}.y_2 + k_{3t}.y_3 - k_{2d}.y_2 \tag{12}$$

$$\frac{dy_3}{dt} = k_{2t}.y_2 - k_{3t}.y_3 - k_{3d}.y_3 \tag{13}$$

$$\frac{dy_4}{dt} = v_{sb}.(\frac{k_{ib}^m}{k_{ib}^m + y_{10}^m}) - k_{4d}.y_4 \tag{14}$$

$$\frac{dy_5}{dt} = k_{4b}.y_4 - k_{5t}.y_5 + k_{6t}.y_6 - k_{5d}.y_5 \tag{15}$$

$$\frac{dy_6}{dt} = k_{5t}.y_5 - k_{6t}.y_6 + k_{7a}.y_7 - k_{6a}.y_6 - k_{6d}.y_6 \tag{16}$$

$$\frac{dy_7}{dt} = k_{6a}.y_6 - k_{7a}.y_7 - k_7.y_7.y_3 - k_{7d}.y_7 \tag{17}$$

$$\frac{dy_8}{dt} = v_{sr}.(\frac{y_7^h}{k_{ar}^h + y_7^h}) - k_{8d}.y_8 \tag{18}$$

$$\frac{dy_9}{dt} = k_{9b}.y_8 - k_{9t}.y_9 + k_{10t}.y_{10} - k_{9d}.y_9 - C_2.v_{cdk1}.z_8.(\frac{y_9}{k_p + y_9}) \tag{19}$$

$$\frac{dy_{10}}{dt} = k_{9t}.y_9 - k_{10t}.y_{10} - k_{10d}.y_{10} - C_2.v_{cdk1}.z_8.(\frac{y_9}{k_p + y_9}) \quad (20)$$

In order to achieve an autonomous circadian clock period (TCR) of 24 hrs, we multiply the 10 ODEs representing the circadian clock gene network by a factor of 1.45 and the autonomous cell cycle period (TCC) is imposed as described in Section 1. For the forward coupled system the TCC = 15 hrs and for the reverse coupled case TCC = 16 hrs.

| Parameter Symbol | Numerical Value |
|---|---|
| $v_{sp}$ | $1.5nMh^{-1}$ |
| $k_{ap}$ | $0.7nM$ |
| $k_{1d}$ | $0.7h^{-1}$ |
| $n$ | 2 |
| $k_{2b}$ | $0.3nM^{-1}h^{-1}$ |
| $q$ | 2 |
| $k_{2d}$ | $0.05h^{-1}$ |
| $k_{2t}$ | $0.24h^{-1}$ |
| $k_{3t}$ | $0.02h^{-1}$ |
| $k_{3d}$ | $0.12h^{-1}$ |
| $v_{sb}$ | $1.8nMh^{-1}$ |
| $m$ | 2 |
| $k_{ib}$ | $2.2nM$ |
| $k_{4d}$ | $0.4h^{-1}$ |
| $k_{5b}$ | $0.24h^{-1}$ |
| $k_{5d}$ | $0.06h^{-1}$ |
| $k_{5t}$ | $0.45h^{-1}$ |
| $k_{6t}$ | $0.06h^{-1}$ |
| $k_{6d}$ | $0.12h^{-1}$ |
| $k_{7a}$ | $0.003h^{-1}$ |
| $k_{6a}$ | $0.09h^{-1}$ |
| $k_{7d}$ | $0.09h^{-1}$ |
| $k_7$ | $1nM^{-1}h^{-1}$ |
| $v_{sr}$ | $1.6nMh^{-1}$ |
| $k_{ar}$ | $0.6nM$ |
| $h$ | 2 |
| $k_{8d}$ | $0.2h^{-1}$ |
| $k_{sr}$ | $1.7h^{-1}$ |
| $k_{9t}$ | $0.8h^{-1}$ |
| $k_{10t}$ | $0.4h^{-1}$ |
| $k_{9d}$ | $0.2h^{-1}$ |
| $k_{10d}$ | $0.2h^{-1}$ |
| $v_{cdk1}$ | $1h^{-1}$ |
| $k_p$ | $1.006nM$ |

Table S5: Circadian clock ODE parameter values for Model 2.

### 3 Numerical Integration of the system of ODEs to generate time series

The system of ODEs that represent the coupled circadian clock-cell cycle gene network for both Model 1 and Model 2 can be evolved to generate time series data representing the change in concentration of various proteins and mRNAs. We numerically integrate the ODEs, using the Euler method which states that a differential equation of the form,

$$\frac{dx}{dt} = f(x)$$

can be discretized as follows:

$$\frac{x(t+\Delta t) - x(t)}{\Delta t} = f(x) \tag{21}$$

The ODEs are deterministic in nature, however owing to the inherent stochasticity in the biological system we incorporate noise into this model to generate stochastic differential equations (SDEs) via the Chemical Langevin Equation framework, integrated using the Euler-Maruyama scheme [2]:.

$$x(t+\Delta t) = x(t) + f(x)\Delta t + A\sqrt{\Delta t}N(0,1) \tag{22}$$

where the term $A.N(0,1).\sqrt{\Delta t}$ is the Gaussian noise term added. Here A is the noise coefficient that is set by comparing Inter-mitotic times distribution as described in section 5, and $N(0,1)$ is a Gaussian Random variable with mean 0 and standard deviation 1.

### 4 Fourier Analysis of time series data

To study the phenomenon of entrainment, we utilised Fourier Transforms to generate the spectral density *versus* frequency curves. This allowed us to detect the distribution of frequencies that constitute the oscillatory time series data for the network (say cell cycle) that is being entrained by the other network (say circadian clock) and vice versa. We fed the time series after eliminating the initial transients into the `spectrum` function in the R programming language with parameter values as follows: span = 5, log = "no", plot = FALSE. The default frequency axis for this function is in cycles per sampling interval. It is more intuitive to express the frequency axis in cycles per unit time hence we divide the extracted frequencies by our sampling rate. We also multiplied the spectral density by 2 so that the area under the periodogram equals the variance of the time series[5]. Thus for time series data say X, we run the following block of code to generate the spectral density curves mentioned in the main text (Figure 2) and SI (Figures S2 and S3). These curves demonstrate the phenomenon of entrainment where increasing coupling strengths changes the period of an oscillator closer to the period of the oscillator entraining it.

```
fourier_x <- spectrum(X,log="no",span=5,plot=FALSE)
freq_x <- fourier_x$freq/sampling_rate
spectral_x <- 2*fourier_x$spec
plot(freq_x,spectral_x,xlab="frequency",ylab="spectral density",type="l")
```

### 5 Simulation to generate cellular lineages

The system of SDEs for either model was used to develop a simulation that creates cellular lineages where the progression through cell cycle is mediated by the different protein concentrations. We utilised the IGRAPH package in R to represent the growing population as a set of directed graphs [6, 7]. After initialisation of a set of ancestral cells which formed the base nodes of the graph, we assigned starting concentrations of the molecular species to each ancestral cell. We then numerically integrated the SDEs separately for each cell in order to determine the concentrations in the next time steps. Depending upon concentration of Mitotis Promoting factor(MPF), the cells traverse the different cell cycle phases and the cells that ultimately reach the M-phase divide to give two daughter cells. The thresholds of MPF concentration for the different transitions used in the simulation are as follows:

| Transition | Threshold |
|---|---|
| G1 to S/G2 | [MPF] $\geq$ 0.09nM |
| G2 to M | [MPF] $>$ [Wee1] |
| M to G1 (division) | [MPF] $\leq$ 0.06nM |

Table S6: MPF concentration threshold used in lineage simulation

Upon reaching the M-phase, we remove the mother cell from the population and add two new nodes (representing two daughter cells) to the lineage graph. The daughter cells enter the G1 phase with concentrations of all species identical to that of the mother cell and the simulation runs till a user-defined final number of cells in the population are reached. The lineage simulation is schematised in Figure S1.

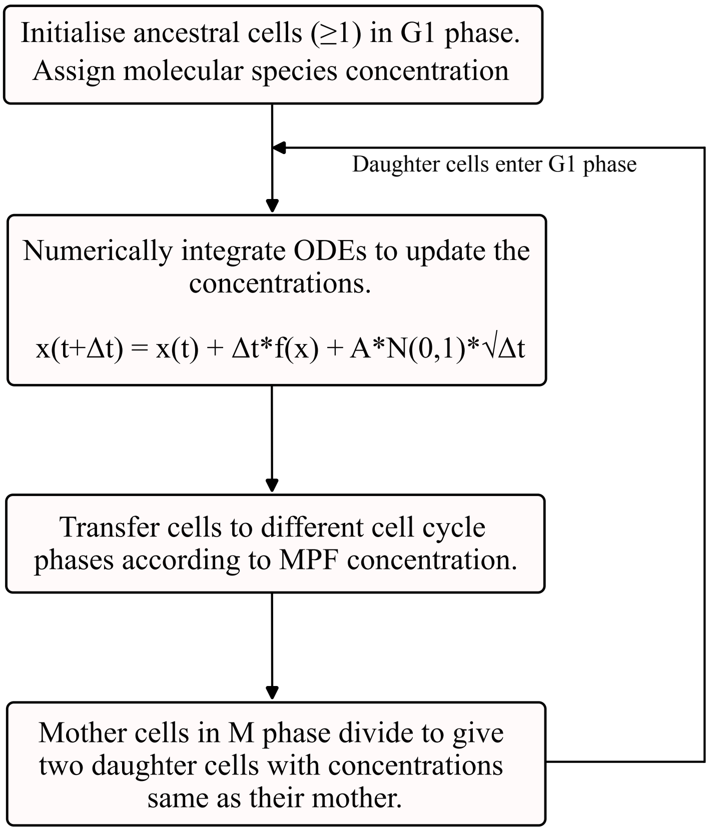


Figure S1: Outline of lineage algorithm

A single run of this simulation generates lineage trees for the starting ancestor cells. From these directed lineage graphs we extracted the inter-mitotic times of each cell as (`division_time -`

birth_time). The noise parameter $A$ in the SDEs of either model (Equation 22) is optimized by matching the simulated IMT distribution with experimental data on HCT116 cells [8]. This parameter was thus set to be 0.025 for Model 1 and 0.003 for Model 2. We extracted unique related cell pairs like sisters, cousins, mother-daughter pairs and determined the Pearson correlation for the IMTs of each pair. Thus one lineage simulation generated a set of correlation values for each type of pair considered indicated by $\rho_{SS}$ for sister correlation, $\rho_{CC}$ for cousin correlation and $\rho_{MD}$ for Mother-Daughter correlation respectively. For each run we also obtained the proliferation rate of the population by fitting a linear model to the $\log(cell\_number)$ versus time plot using the `fit.lm` function in R, after excluding the initial transients. The slope of the fit is the population proliferation rate.

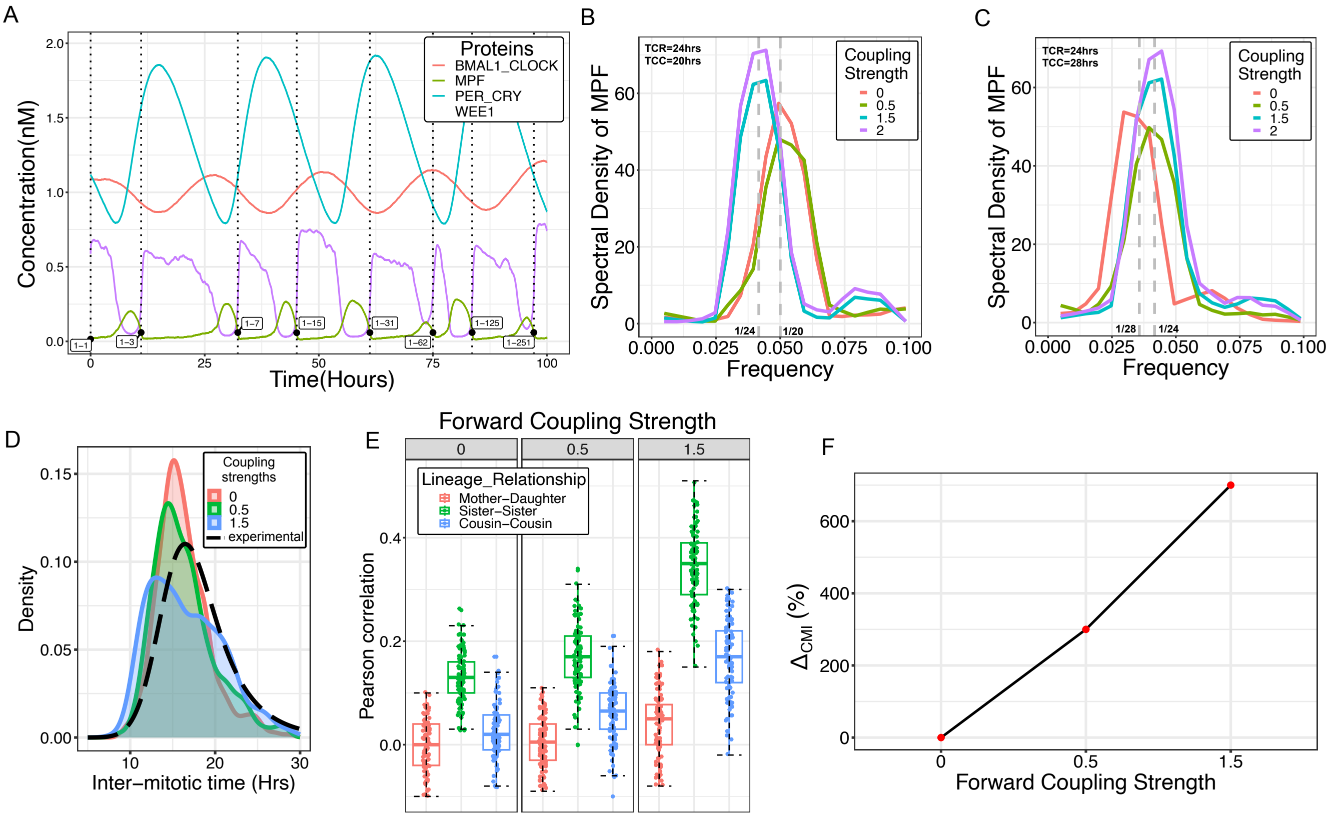


Figure S2: Forward coupled circadian clock – cell cycle gene network correlates Inter-mitotic times in cellular lineages for Model 1: (A) Time evolution of different proteins for a lineage, recapitulating the expected oscillations. The cells belonging to a lineage that were tracked to generate these oscillatory trajectories are labelled at the time of their birth on the graph. The different proteins present in the network satisfy the experimentally observed phase relationships. (B) & (C) Entrainment of cell cycle by the circadian clock. Increasing forward coupling strength changes the cell cycle frequency from 1/20 or 1/28 hour$^{-1}$ respectively to the circadian clock frequency of 1/24 hour$^{-1}$. (D) With increasing forward coupling strength the distribution of the Inter-Mitotic times (demonstrated by the different colours) match the experimentally observed distribution. (E) The emergence of high sister correlation and also cousin-mother inequality with increasing coupling strength as seen in Model 2. (F) Percentage change in the difference between the median value of cousins and mother-daughter correlation (i.e. cousin-mother inequality) in comparison to the uncoupled system, shown for the different coupling strengths considered for the forward case. (Boxplot and median value calculated for 100 runs of simulation. Here TCC=16hrs)

For a fixed parameter set (denoted by $i$), We ran a set of 100 simulations and generated a distribution of Pearson correlation values for sisters ($\rho_{SS}$), cousins ($\rho_{CC}$) and mother-daughter pairs ($\rho_{MD}$). From this distribution we calculated the cousin-mother inequality for this parameter set as follows:

$$\rho^i_{CMI} = \text{median}(\rho_{CC}) - \text{median}(\rho_{MD})$$

We calculated $\rho^i_{CMI}$ for different parameter sets (for example by changing the coupling strength or KL001 concentration) and then calculated the percentage change in cousin-mother inequality for each parameter set $i$ compared to a control parameter set, $\Delta^i_{CMI}$:

$$\Delta^i_{CMI}\% = \frac{(\rho^{(i)}_{CMI} - \rho^{(control)}_{CMI})}{\rho^{(control)}_{CMI}} * 100$$

Therefore when $i$ represents the control set of parameters, $\Delta^i_{CMI}\%$ is 0. In case of simulations with different coupling strengths, similar to the behaviour of Model 2 (main text Figure 2), we observed emergence of the cousin-mother inequality for increasing coupling strengths in Model 1 as well (Figure S2).

For each of the 100 simulations that we ran for a particular parameter set, we also calculated the proliferation rate of the population to generate a distribution of proliferation rates. The percentage change in proliferation rate $\Delta_{growth}\%$ with respect to a control set of parameters is then similarly calculated as for the cousin-mother inequality described above. If the median population growth rate for the 100 runs of the simulation for the $i^{th}$ parameter set is indicated as $GR^{(i)}$ then $\Delta_{growth}\%$ is calculated as:

$$\Delta_{growth}\% = \frac{(GR^{(i)} - GR^{(control)})}{GR^{(control)}} * 100.$$

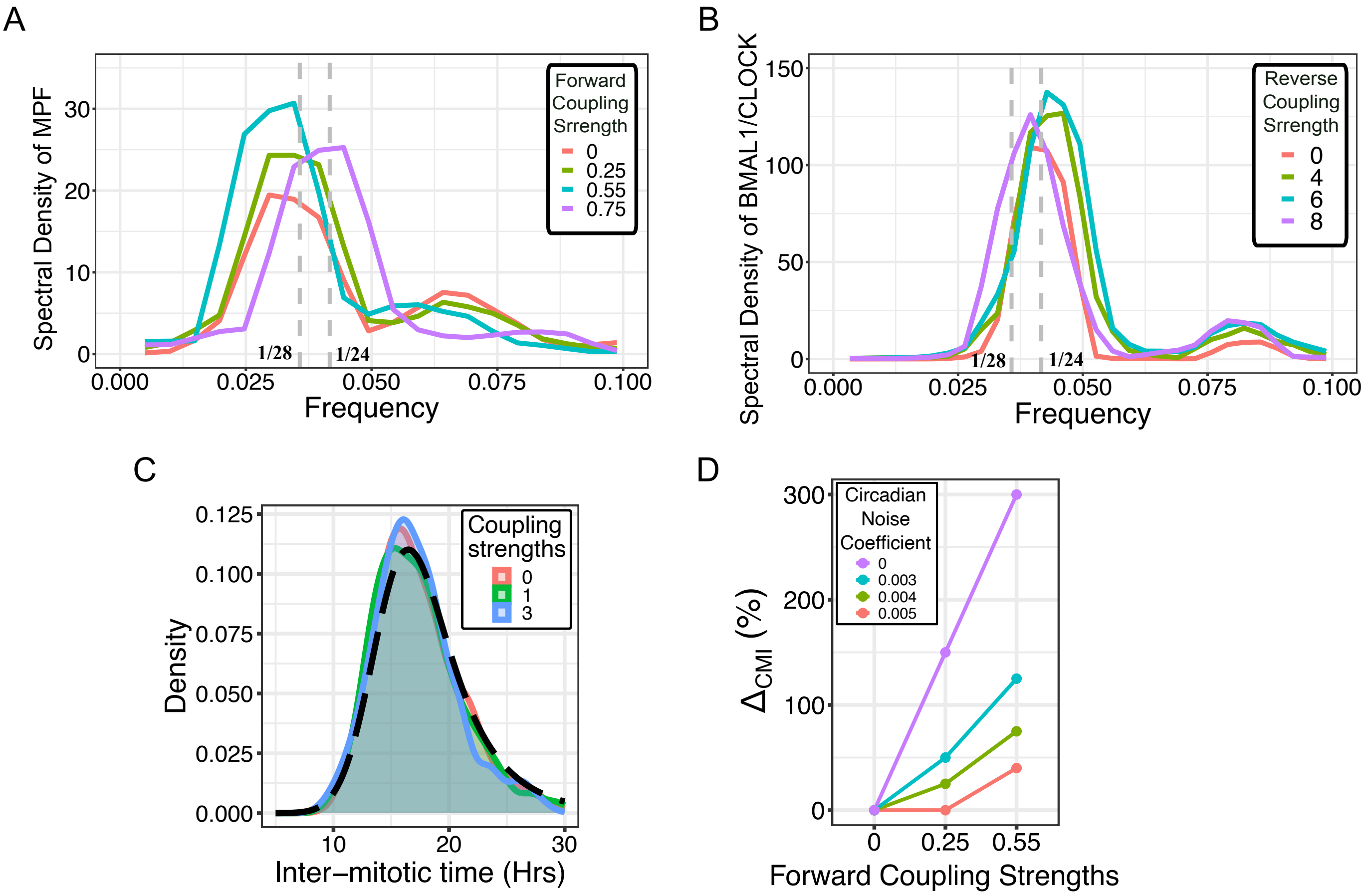


Figure S3: Supplementary results for Model 2: (A) Entrainment of the cell cycle by circadian clock. With increasing forward coupling strength the frequency of oscillation of cell cycle gene MPF changes from 1/28 to 1/24 hour$^{-1}$. (B) Entrainment of circadian clock by the cell cycle. With increasing reverse coupling strength the circa- dian clock heterodimer BMAL1/CLOCK oscillation frequency changes from 1/24 to 1/28 hour$^{-1}$. Circadian clock period (TCR)=24 hrs, Autonomous cell cycle period (TCC)=28 hrs. (B) Comparison of IMT distribution from simulation and experiment for a reverse coupled system. The simulated IMT distribution matches the experimentally observed IMT distribution (black dashed line). (D) Effect of incorporating different noise coefficients for the circadian clock while keeping the noise coefficient for cell cycle fixed to what is mentioned in Section 5. With increasing noise in the circadian clock gene network the percentage change in cousin mother inequality $\Delta_{CMI}\%$ for C1 $\geq 0$ when compared to uncoupled system that is C1 = C2 = 0, decreases.

### 6 Simulating KL001 mediated circadian clock inhibition

In order to mimic the effect of KL001 mediated circadian clock inhibitor, we decreased the degradation rates of the PER-CRY complex [9]. KL001 prevents ubiquitin - mediated degradation of the CRY proteins. Previous studies have shown that the degradation occurs both in cytoplasm as well as nucleus, hence we divided both the degradation rates $k_{2d}$ and $k_{3d}$ by the numbers mentioned in the graphs in main text Figure 3 as well as SI Figure S4 to incorporate KL001's effect. Dividing by 1 is the control, where the degradation rates are unchanged and hence inhibition is absent. Division by higher numbers reduces the degradation rates, and represents higher KL001 concentrations. The percentage change in cousin-mother inequality and proliferation rate for the different KL001 inhibition cases were determined as mentioned in section 5. For Model 1 we observed a decrease in the $\Delta_{CMI}\%$ for higher KL001 values ($> 1$) similar to the behaviour observed for Model 2 in main text Figure 3. However unlike in Model 2, where the change in $\Delta_{growth}\%$ was minimal, in case of Model

1 the behaviour was erratic and no specific trend was observed (Figure S4).

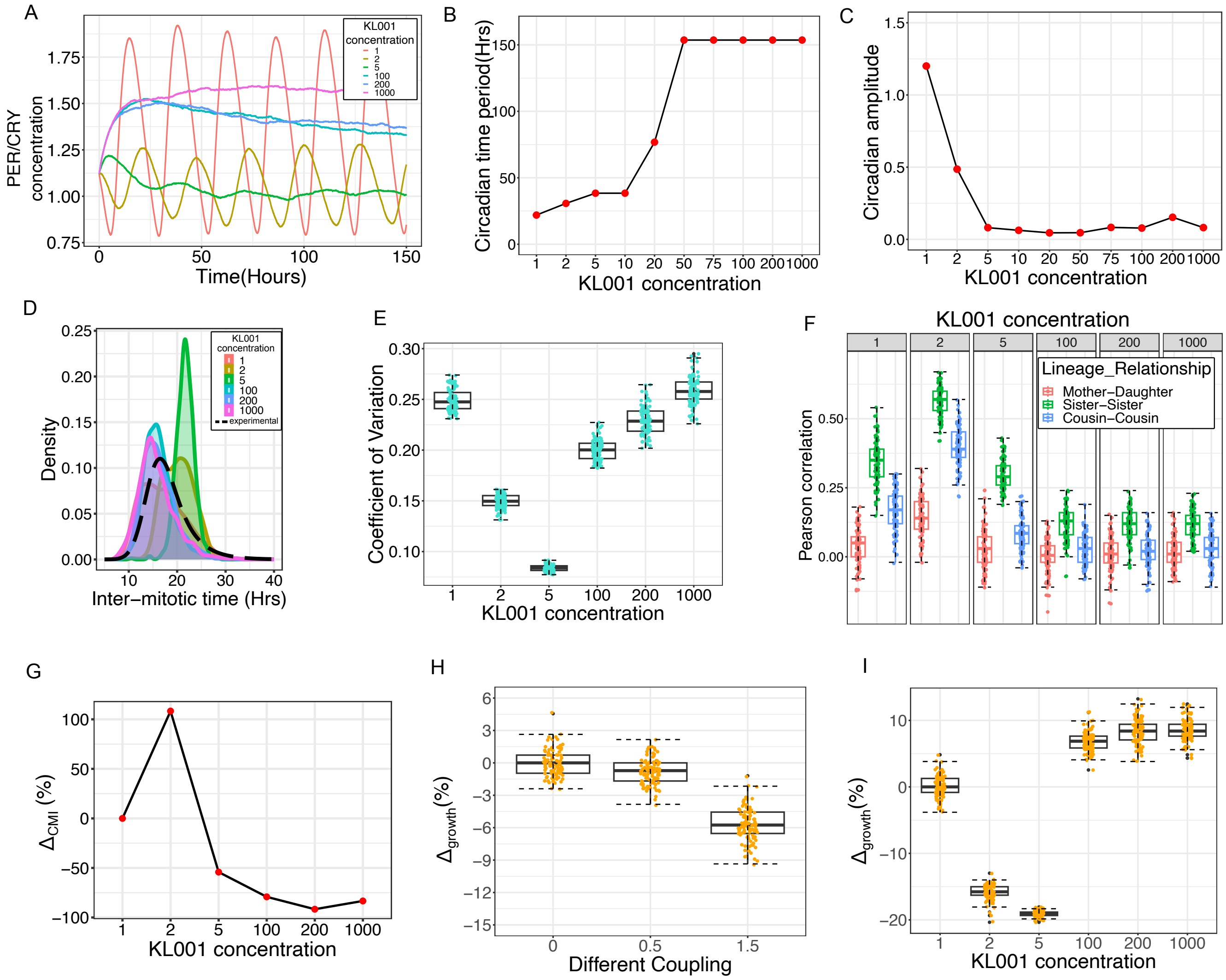


Figure S4: Circadian clock perturbation with KL001 decreases IMT correlations for Model 1: (A) Damped circadian proteins' oscillations as observed for increasing concentrations of KL001. (B) & (C) Change in time period and amplitude of circadian clock as observed under influence of KL001 (D) Changes in IMT distribution for different KL001 concentration. (E) Effect of KL001 on the Coefficient of variation of IMT distribution. While for Model 2 we observed a decrease in the variance of IMT distribution with increasing KL001, we do not observe any such trend for Model 1. (F) KL001 diminishes the cousin-mother inequality in lineage simulation similar to behviour observed for Model 2. (G) Percentage change in median value of cousin-mother inequality in comparison to the control case, shown for the different concentrations of KL001. (H) Change in population growth rate for increasing coupling strengths, predict a decrease due to increase in the cell cycle period owning to entrainment by the circadian clock. (I) Change in growth rate when KL001 addition is simulated the change in growth rate is erratic and no visible trend is present. (Boxplot and median value calculated for 100 runs of simulation. Here C = 1.5, TCC=16hrs, for the lineage simulations under KL001 effect.)

### 7 Digital extraction of KL001 data and downstream processing

We obtained the luminescence rhythm data of Bmal1-dLuc and Per2-dLuc reporter U2OS, Human osteosarcoma cell line under the influence of different concentrations of KL001 from published literature [9]. Using publicly available software DataThief III [10], we digitally extracted the data points from the graphs. Having obtained a time series, we performed Fourier Transform as defined earlier in Section 4 to extract the time period (1/frequency with maximum spectral density) of oscillations for the different concentrations of KL001. Eliminating the initial transients, we also extracted the average amplitude over the different oscillatory cycles.
